# HadoopCNV: A dynamic programming imputation algorithm to detect copy number variants from sequencing data

**DOI:** 10.1101/124339

**Authors:** Hui Yang, Gary Chen, Leandro Lima, Han Fang, Laura Jimenez, Mingyao Li, Gholson J Lyon, Max He, Kai Wang

## Abstract

**BACKGROUND:** Whole-genome sequencing (WGS) data may be used to identify copy number variations (CNVs). Existing CNV detection methods mostly rely on read depth or alignment characteristics (paired-end distance and split reads) to infer gains/losses, while neglecting allelic intensity ratios and cannot quantify copy numbers. Additionally, most CNV callers are not scalable to handle a large number of WGS samples.

**METHODS:** To facilitate large-scale and rapid CNV detection from WGS data, we developed a Dynamic Programming Imputation (DPI) based algorithm called HadoopCNV, which infers copy number changes through both allelic frequency and read depth information. Our implementation is built on the Hadoop framework, enabling multiple compute nodes to work in parallel.

**RESULTS:** Compared to two widely used tools – CNVnator and LUMPY, HadoopCNV has similar or better performance on both simulated data sets and real data on the NA12878 individual. Additionally, analysis on a 10-member pedigree showed that HadoopCNV has a Mendelian precision that is similar or better than other tools. Furthermore, HadoopCNV can accurately infer loss of heterozygosity (LOH), while other tools cannot. HadoopCNV requires only 1.6 hours for a human genome with 30X coverage, on a 32-node cluster, with a linear relationship between speed improvement and the number of nodes. We further developed a method to combine HadoopCNV and LUMPY result, and demonstrated that the combination resulted in better performance than any individual tools.

**CONCLUSIONS:** The combination of high-resolution, allele-specific read depth from WGS data and Hadoop framework can result in efficient and accurate detection of CNVs.

## INTRODUCTION

Copy number variation (CNV) refers to copy number changes of a DNA fragment usually between one kilobases (kb) and less than five megabases (Mb) (1). CNV is one of the major forms of genomic variations. Based on published high-quality data on healthy individuals of various ethnicities, it is estimated that 4.8–9.5% of the genome contributes to CNVs (2). Numerous studies have reported that CNVs are associated with a large number of human diseases (3-5). Before the next-generation sequencing (NGS) era, most CNVs documented in literature were identified through microarrays, with help from software tools such as PennCNV (6), QuantiSNP (7) and many others (8-11). Although CNV discovery with microarrays was successful in discovering a large number of disease associated CNVs, the limited resolution of microarrays is not capable to identify all CNVs, therefore a large number of disease-contributing CNVs might have been overlooked.

NGS technologies are increasingly used to find disease-contributing variations. The high-throughput capacity of NGS makes it practical to identify most CNVs in the genome that were difficult to be discovered previously. Many pioneering studies have demonstrated the feasibility of discovering CNVs through the use of NGS data, by examining the alignment files directly. Several approaches were developed and each had different advantages and disadvantages (12). For small-scale CNVs, paired-end mapping and split-read are useful for detection, and tools such as BreakDancer (13), AGE (14), GASV (15), PINDEL (16), LUMPY (17), DELLY (18) were developed and widely used today. For larger CNVs, read-depth based approaches such as CNV-Seq (19), BIC-Seq (20), JointSLM (21) and CNVnator (22) have been demonstrated to be more useful. Among them, CNVnator (22) uses a mean-shift algorithm to identify the boundaries of CNVs and proceeds with multiple refinement and GC correction, demonstrating the clear utility of read depth for CNV calling. Assembly based approaches have also been useful for CNV detection, such as the algorithms used by Cortex assembler (23) and TIGRA-SV (24), but these approaches are typically not computationally efficient to be used in genome-wide analysis. Finally, several tools were developed specifically for CNV detection from exome sequencing data, such as XHMM (25), CODEX (26), ExomeCNV (27), ExomeCopy (28) and ExCopyDepth (29).

Hadoop framework has been demonstrated to be able to handle large-scale sequence data (30). Here we present a scalable CNV calling algorithm within a *big data* framework, HadoopCNV, which rapidly extracts read depth information from sequence alignment (BAM) files. HadoopCNV integrates three components into a MapReduce based pipeline that seamlessly runs on environments ranging from a single computer to thousands of computers. The first component measures depths for each allele at user-specified intervals. The second component smoothes these measurements by defining bins of a user-specified dimension (100 bp by default), and storing summary statistics at each bin. The final component invokes a Viterbi scoring algorithm using the model previously described (31). HadoopCNV leverages the Hadoop Distributed File System (HDFS), making it possible to store a large number of samples and call CNVs on them in a distributed manner among tens, hundreds or even thousands of nodes, thus provides a *big data* solution to genome analysis. Considering that some large-scale genome centers are migrating to Hadoop to store alignment (BAM) files, and that several computational tools are already developed to perform genome analysis on BAM files (32), the availability of HadoopCNV therefore also allows seamless integration into existing analytical pipelines.

## RESULTS

### Overview of the approach

The basic framework for HadoopCNV is described in **Figure 1**. Since the size of whole genome sequencing (WGS) data is generally large, it is critical to have a scalable infrastructure to generate CNV calls, especially when a large number of samples need to be processed in a short period of time. We adopted Apache Hadoop (http://hadoop.apache.org/), which is an open-source infrastructure that allows for reliable and scalable distributed processing of very large data sets across clusters built from multiple computers using HDFS (Hadoop Distributed File System), YARN (Yet Another Resource Negotiator) and the MapReduce programming models. HDFS is a Java-based file system that provides scalable and reliable data storage, and it was designed to span large clusters of commodity servers. YARN is a framework for job scheduling and cluster resource management, such as managing and monitoring workloads, maintaining a multi-tenant environment, implementing security controls, and managing high availability features of Hadoop. HadoopCNV takes BAM and VCF files of a WGS data as input. The BAM file needs to be available from HDFS while the relevant VCF file can be located anywhere at a user’s local file system. The CNV calling algorithm is then executed on multiple machines simultaneously with the MapReduce paradigm. The output from HadoopCNV is a list of regions, each annotated with deletion or duplication and the corresponding copy numbers. HadoopCNV is freely available at http://www.github.com/WGLab/HadoopCNV.

**Figure 1.**
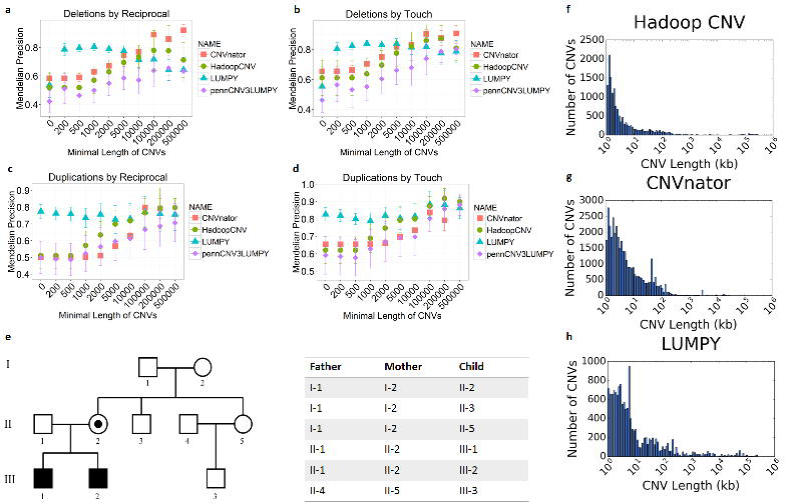
Overview of HadoopCNV. All the BAM files are stored in Hadoop File System and the VCF file can be stored in the local hard drive. 1) Alignments from each BAM file are first extracted, then generated for each bin in each chromosome with the Depth Caller MapReduce. 2) With the reads for each bin at each chromosome are aggregated, then with the help of VCF information, the median depth and SSMSE are extracted for each bin with the Region Binner MapReduce. Finally, the Median Depth and SSMSE are aggregated for each chromosome first, then the copy number for each position range is calculated, which is the final output from the software.

### Speed comparison with other CNV calling methods

To illustrate the speed advantage of HadoopCNV, we compared the running time among LUMPY (33), CNVnator (22) and HadoopCNV on a BAM file with ~30X coverage for NA12878, which is generated from the 1000 Genome Project. We recorded the timestamps before the CNV calling program was executed and after the program was finished. Since the performance of any Hadoop system depends on how many data nodes it has, we measured the running time of HadoopCNV on a cluster built with 2 data nodes, 6 data nodes, 10 data nodes, 16 data nodes, and 32 data nodes, respectively. With 32 data nodes, HadoopCNV only takes 1.6 hours to process one whole-genome, which is about one sixth the time used by CNVnator (**Figure 2**). Meanwhile, we confirmed that the speed approximately linearly scales with the number of data nodes in the Hadoop cluster, demonstrating the efficacy of our data-parallel algorithm.

**Figure 2.**
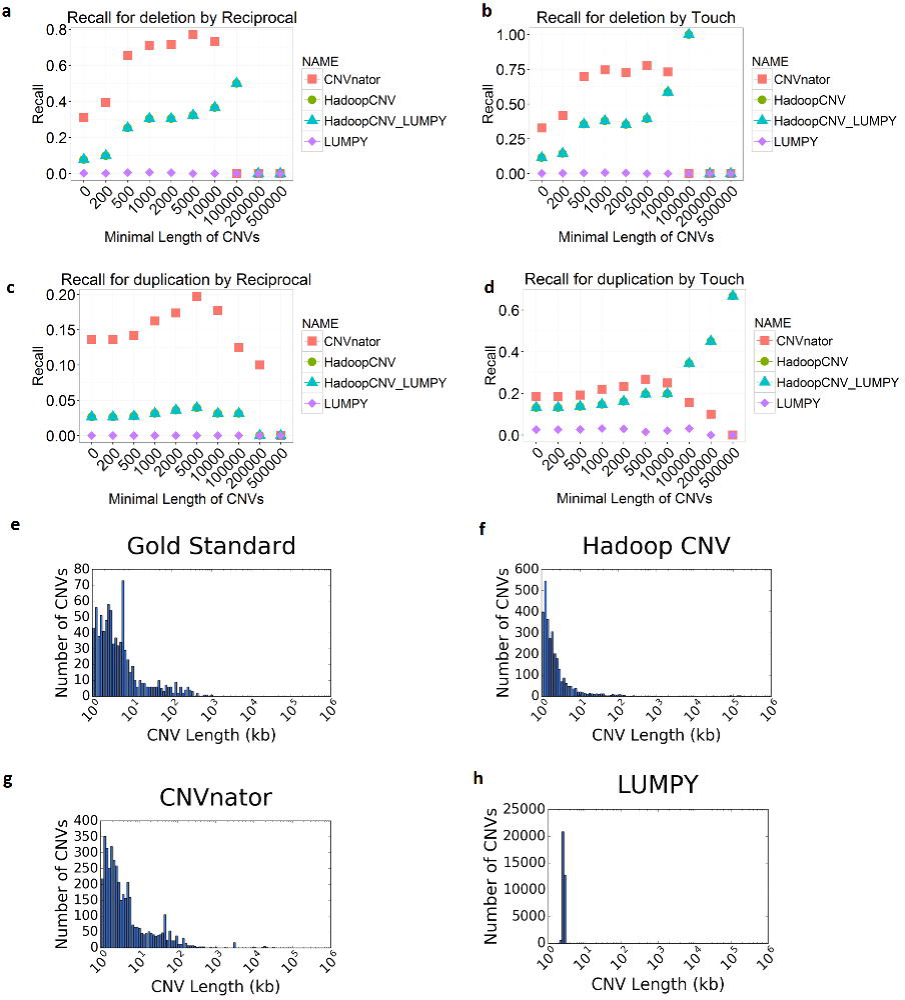
Speed comparison for HadoopCNV with 2, 6, 10, 16, 32 nodes, CNVnator and LUMPY. HadoopCNV only costs about 1.6 hours for a WGS data with 32 nodes. The number of replicate experiment is 3 for HadoopCNV, and 5 for CNVnator and LUMPY. Error bars represent standard deviation.

### Evaluation on simulated datasets

To evaluate the accuracy of different CNV callers including HadoopCNV, we simulated 20 deletions and 20 duplications with different sizes in each chromosome (480 CNVs in total), and then simulated individual genomes with these CNVs (see Methods for details). From the genome sequence, we then simulated sequencing reads, which consist of 100-bp reads as FASTQ files with 40X genome coverage. We mapped these reads to the human genome (build hg19) by BWA (34), generated BAM files and then tested HadoopCNV, CNVnator and LUMPY on these files. This approach allows us to evaluate their performance based on their concordance with the ground truth, and assess how performance changes as the minimal CNV length threshold increases. We assessed two types of “concordance” measures, touch (any overlap) and 50% reciprocal overlap (see Methods for details). For deletions, HadoopCNV has a consistently higher concordance with ground truth, compared to CNVnator by both standards, and a similar performance as LUMPY. For duplications, HadoopCNV has similar performance with other tools, especially when the length threshold is larger than 15k, we observed a consistent performance advantage of Hadoop CNV over other tools (**Figure 3f-i**). In addition, with our approach of combining HadoopCNV and LUMPY, we are able to achieve a better performance than any other individual tool in any scenario. Also, from the precision and recall comparison, we can observe that CNVnator performs relatively worse at precision, but HadoopCNV has a lower recall when the threshold is small **(Supplemental Figure 1).** The length distribution of the CNV calls shows that these tools are largely comparable with each other (**Figure 3a-b**).

**Figure 3.**
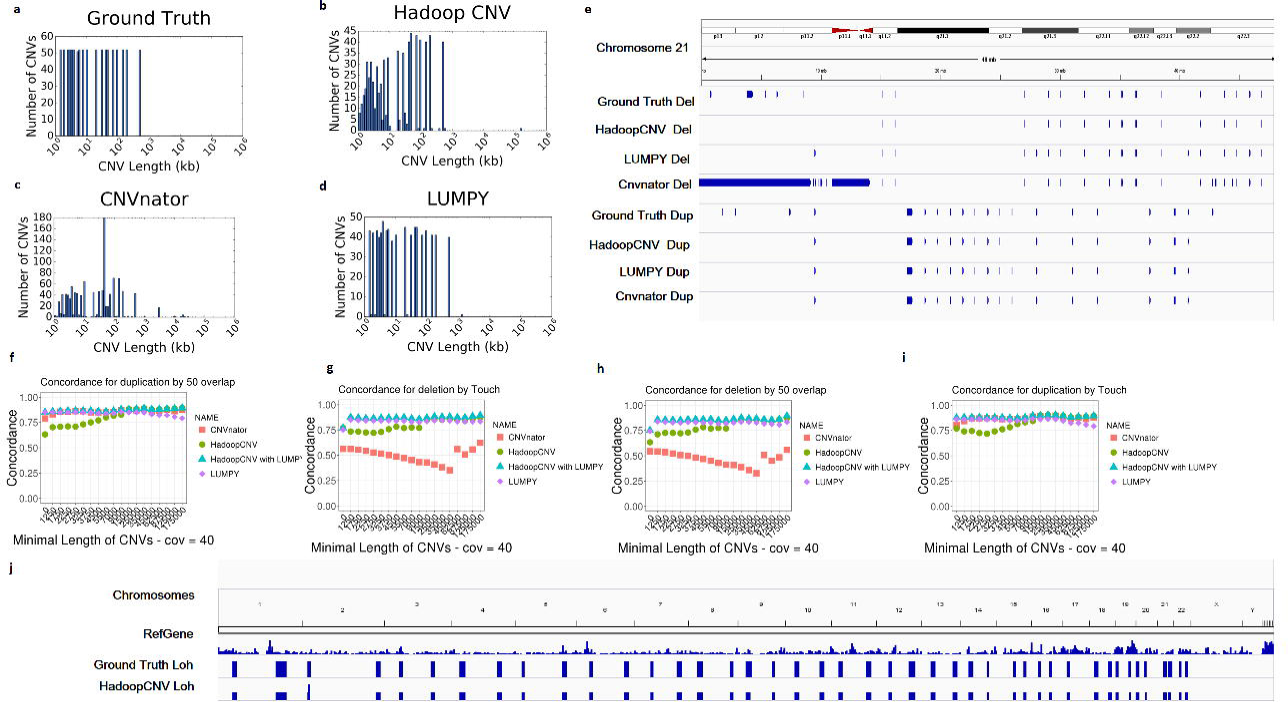
Comparison of HadoopCNV and CNVnator on the simulated samples with CNVs. The length distribution of the ground truth CNV to generate simulations (a), HadoopCNV output (b), CNVnator output (c) and LUMPY output (d). A snapshot of all the deletions and duplications in chromosome 21 (e). The concordance between the output CNVs and the ground truth at different length thresholds were shown, for deletions with 50% reciprocal standard (f) and touch standard (g), and for duplications with 50% reciprocal standard (h) and touch standard (i). A snapshot of Ground Truth LOHs and the ones called by HadoopCNV is also shown (j).

We also generated an IGV (Integrative Genome Viewer) snapshot of all the CNVs located on chromosome 21. From the snapshot, we noticed that CNVnator made a very large deletion call from this IGV snapshot. After further investigation, we found that this comes from a region that has all Ns in the reference genome. Therefore, it is important to post-process CNV calls from CNVnatore. In contrast, HadoopCNV and LUMPY successfully skipped this region (**Figure 3e**).

### Detection of copy-neutral loss-of-heterozygosity (LOH)

Due to its unique use of allelic information in sequence alignments, HadoopCNV is able to identify copy-neutral loss of heterozygosity (LOH) in the genomes, by treating it as a separate state in the model. Large LOH events typically signify genomic segments that are identity by descent (for example, due to consanguinity), or created by uniparental disomy, and may be of clinical significance, especially when the LOH region harbors disease variants for recessive diseases (35). In comparison, other tools such as CNVnator and LUMPY cannot identify LOH in their output.

To demonstrate the ability of HadoopCNV to identify copy-neutral LOHs, we simulated two LOHs on each chromosome of varying sizes (See Methods). From the result, HadoopCNV is able to identify all these regions, although calling two LOHs instead of one in one case in chromosome 4 (**Figure 3j**). The overall concordance based on 50% reciprocal overlap and touch standards are 91.3% and 95.6%, respectively. In summary, through the use of allelic information, HadoopCNV is capable to identifying copy-neutral LOH events, which may be missed by other CNV callers that ignore such information.

### Evaluation on the NA12878 sample

To evaluate the performance of HadoopCNV on real data, we first used the NA12878 sample from the 1000 Genomes Project for comparison, which was generated by Illumina HiSeq. We used this sample because it has been comprehensively studied by multiple previous CNV studies (36). Moreover, a “gold standard” dataset has been previously compiled from NA12878 (37). For deletions, we updated the gold standard with a latest benchmark dataset (38). However, we acknowledge that although this ‘gold standard’ set is widely used in literature, it might miss many true CNV events as most of the CNVs in the gold standard are required to be verified by multiple calling methods, some of which have low resolutions. Thus here we focused on a recall comparison, to evaluate how many gold standard CNVs can be captured by different methods.

We compared our method with LUMPY and CNVnator, which are among the most widely used CNV calling methods for WGS data. Our results indicate that CNVnator generally has a slightly better recall. HadoopCNV is in general better than LUMPY, for both deletion and duplication. HadoopCNV has a consistently better performance than LUMPY (**Figure 4a-d**). The combination between HadoopCNV and LUMPY is compared here as more CNVs definitely mean a better recall, different from concordance comparison. The CNV length distributions are also shown for the gold standard and the outputs from these three tools, where LUMPY has a relatively narrow distribution, due its unique use of split read and read pair (**Figure 4e-h**). Combining the results from both the simulation data and NA12878 data, we conclude that HadoopCNV has comparable performance with the commonly used read depth based CNV calling tool CNVnator, and even better performance for large CNVs.

**Figure 4.**
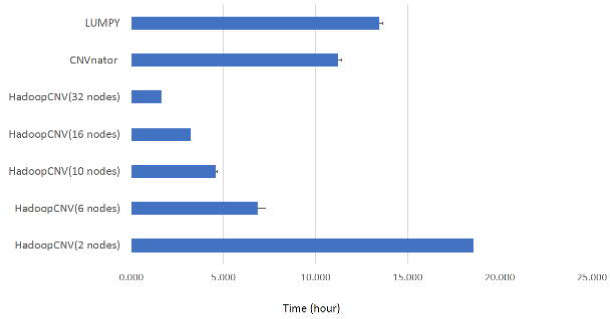
The recall comparison for gold standard CNV calls of NA12878, among HadoopCNV, CNVnator and LUMPY, calculated by averaging the results of three different NA12878 sequencing samples. The recall comparison for deletion with 50% reciprocal overlap standard (a) or touch standard (b), and for duplication with 50% reciprocal standard (c) and (d) is shown. The CNV length distribution is shown as histograms, for gold standard (i), HadoopCNV output (j), CNVnator output (k) and LUMPY output (j). For the purpose of a better display, the CNVs larger than 100 kb are categorized into 100 kb bin. Error bars represent standard deviation within the 3 samples.

### Comparison of CNV calling accuracy on a 10-member pedigree

Since NA12878 has been used by almost all CNV publications to date, thus it is likely that many other software tools are already tuned for NA12878 to achieve good performance by comparing to the published ‘gold standard’. Therefore, it is important to have a different dataset for unbiased evaluation of the performance of different CNV callers. Additionally, since the vast majority of CNVs are inherited, Mendelian precision is a reliable measure of precision of CNV calling algorithms, which is the percentage of CNVs in the child that are also found in parents. To further compare the performance of HadoopCNV against other existing software programs, we analyzed a WGS dataset generated from a 10-member pedigree, which was previously sequenced at ~50X coverage per individual. We used this pedigree to compare Mendelian precisions among different CNV callers, as measures of false positive/negative rates for CNV calls.

To evaluate Mendelian precisions, we selected all six trios in the pedigree (**Figure 5e**) and calculated the proportion of CNVs that are not detected in parents, based on the 50% reciprocal overlap and touch standards. For each comparison, we filtered CNV calls with different minimal length thresholds. We hypothesized that the higher the length threshold, the more likely that the CNV call is due to false positive in offspring or false negative in parents. We acknowledge the potential existence of *de novo* CNVs; however, given the very low prior probability of having a *de novo* event, we ignore this possibility in this evaluation. We calculated the Mendelian precision by one minus Mendelian error rate and conducted comparisons among HadoopCNV, CNVnator and LUMPY (**Figure 5a-d**). This comparative analysis demonstrated that HadoopCNV is comparable with CNVnator and LUMPY. At low length threshold, LUMPY generally has a slightly better Mendelian precision - for example, at 200bp threshold, LUMPY has 80% precision compared with around 50% precision of HadoopCNV and CNVnator for duplications, based on 50% reciprocal overlap (**Figure 5c**). At high length threshold, LUMPY performs slightly worse than HadoopCNV and CNVnator, probably caused by their different algorithms.

**Figure 5.**
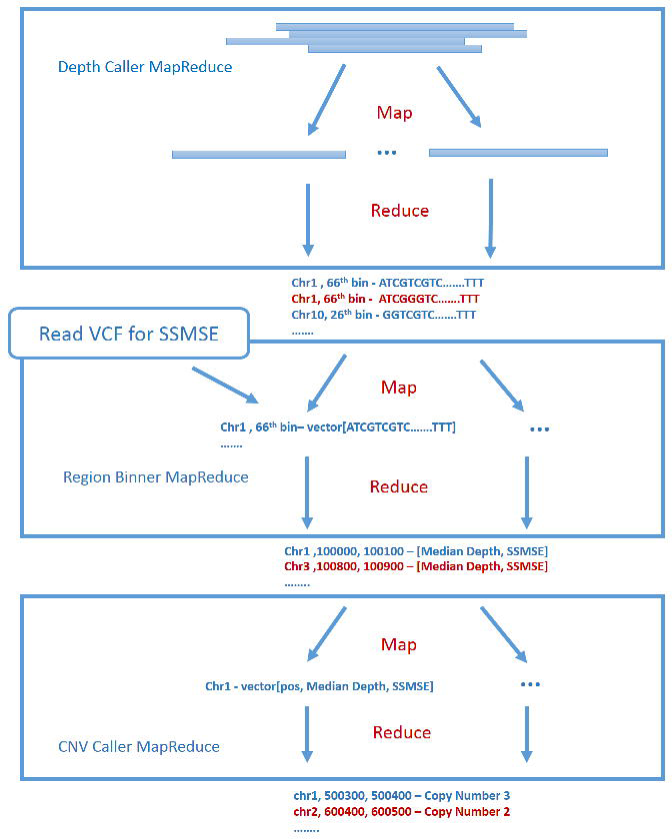
Comparison of the average Mendelian precision (1-Mendelian error rate) based on the 10-member family CNV calls. The Mendelian precision for deletions with 50% reciprocal standard (a) and touch standard (b), and for duplications with 50% reciprocal standard (c) and touch standard (d) are shown. The 10 individual family tree and the 6 trios are shown (e). The CNV length distributions for all CNVs from these 10 individuals are shown, for HadoopCNV (f), CNVnator (g) and LUMPY (h). For the purpose of a better display, the CNVs larger than 100 kb are categorized into 100 kb bin. Error bars represent the standard deviation within the 6 trios.

## DISCUSSION

In this study, we applied the Hadoop framework to implement a highly parallel and scalable CNV calling tool called HadoopCNV. We compared the performance of HadoopCNV with other CNV calling tools on both computational speed and calling accuracy. For speed, HadoopCNV is at least 6 times faster than CNVnator running in a 32-node Hadoop framework. For the comparison on simulated CNVs, HadoopCNV has a slightly better performance than CNVnator, especially for large deletions. For NA12878 gold standard, HadoopCNV has a comparable performance. For Mendelian precision, CNVnator, LUMPY and HadoopCNV have similar performance.

Compared to other CNV calling methods, HadoopCNV has several advantages. First, we utilize a Hadoop framework in our software framework, enabling us to call CNVs in parallel, which substantially speeds up our CNV calling process. From our timestamp evaluation, with 32 data nodes, our CNV caller is at least 6 times faster than CNVnator, which to our knowledge is one of the most commonly used CNV callers for WGS data. Based on our time benchmarking result, the more nodes we have, the faster speed we can achieve from HadoopCNV, and it appears to be linearly scalable. To the best of our knowledge, no other tools are able to use a Hadoop framework for highly parallel CNV calling, thus the advantage of HadoopCNV will be more obvious when handling large-scale datasets such as thousands of whole genomes. Second, we address here the importance of a scalable solution in the *big data* era of human genomics. Similar to CNV calling, other tasks such as SNV calling, indel calling and read alignment and even just data storage are difficult to scale, under current practices in genome analysis (i.e. local file storage, SGE-based parallelization), especially when we face thousands, tens of thousands, or even hundreds of thousands of WGS samples in the future. Thus a highly scalable *big data* framework and standard are urgently needed, and this framework needs to be adopted by the scientific community to prepare for the future. Third, HadoopCNV uses read depth information and alternative allele frequency information, and integrates them into a single coherent model for the most powerful detection of CNVs. To our knowledge, this is among the first computational approaches that integrate the allelic intensity ratio information for CNV calling with WGS data.

HadoopCNV was developed on Apache Hadoop, which is an open-source software infrastructure for distributed, scalable, and reliable computing. One potential improvement is that HadoopCNV can be further speed up by storing variant information from VCF files and read depth of every site on a whole genome from BAM files into a NoSQL database, such as HBase, Cassandra, MongoDB, and others. We have previously built a variant annotation warehouse called SeqHBase (32) that performs similar function in HBase, and we may implement this feature in a future release in HadoopCNV.However, we need to point it out here that although HadoopCNV is able to achieve similar performance as LUMPY, HadoopCNV is not adept at catching small CNVs, especially the ones with a few hundred bps, due to its binning method with 100bp bin, which is the same as CNVnator. Thus we developed a novel method to combine HadoopCNV result and LUMPY result, and demonstrated the combined method has a better performance than any of these compared tools (**Figure 3**). Also, for breakpoint errors, we found HadoopCNV itself performs worse than the other two methods. However, after combination with LUMPY, it can achieve a better breakpoint error than either tool **(Supplemental Figure 1).** We also noticed that combination between LUMPY and HadoopCNV reduces Mendelian precision (**Figure 5**), further suggesting that these two tools are highly complementary to each other.

In addition, we stress here that HadoopCNV is not able to identify other types of structural variations such as inversions and balanced translocations, also due to its read depth method. Another disadvantage of HadoopCNV is the requirement of Hadoop framework adds another layer of technical challenges to biologists and bioinformaticians, which requires basic understanding and skills to operate the YARN and HDFS system.

Finally, we stress that genomic regions with low complexity could be filtered out to help improve CNV qualities. For example, the PennCNV package has instructions on how to post-process and clean CNV calls. However, there are multiple ways of filtration and there are multiple regions that may be filtered, such as immunoglobin regions, centromeric regions, telomeric regions, low complexity regions, and heterochromatin regions. Therefore, we leave options to the users to perform post-processing of the CNV calls.

In summary, HadoopCNV is an efficient and reliable CNV calling program. It demonstrated satisfactory performance on WGS data from a 10-member pedigree and samples from the 1000 Genomes Project. With the rapid decline of sequencing cost for WGS and the continued adoption of WGS in clinical settings, we believe that HadoopCNV will play an important role in genome analysis and can facilitate the implementation of genomic medicine.

## Materials and Methods

### Simulated data set

We used human genome (hg19) as our reference genome for data simulation. We created two independent copies of genome with single-nucleotide variants (SNVs) based on allele frequencies for the AFR population from the 1000 Genomes project (39). Next, we simulated both deletions and duplication, for chromosomes 1 to 22, with sizes 1kb, 1.5kb, 2kb, 2.5kb, 3kb, 3.5kb, 4kb, 5kb, 6kb, 8kb, 10kb, 20kb, 30kb, 40kb, 50kb, 75kb, 100kb, 150kb, 200kb and 500kb, with a minimum distance of 100kb between any two CNVs. These variants were used as ground truth in our comparison. Next, we generated simulated Illumina paired-end short reads with the tool ART(40) and aligned them to the hg19 reference genome using BWA-mem (34). The generated BAM files were used as input for HadoopCNV, CNVnator and LUMPY. The LOHs were simulated based on real LOHs from MCG CNV Database (http://www.cghtmd.jp/CNVDatabase/cnv-summary-map).

### Testing data set (NA12878)

The whole genome sequence (WGS) data for HapMap subject NA12878 were retrieved from multiple resources, because this subject was sequenced multiple times at different institutions: (1) we downloaded the high-coverage (>30X) data from the 1000 Genomes Project site (ftp://ftp.1000genomes.ebi.ac.uk/vol1/ftp/technical/pilot2_high_cov_GRCh37_bams/data/). This is part of the pilot project, where six genomes were sequenced at high coverage. (2) We downloaded the high-coverage (>30X) data generated by HiSeq X Ten from the Garvan Institute (https://dnanexus-rnd.s3.amazonaws.com/NA12878-xten.html).

### Testing data set (10-member pedigree)

We also compared the performance of HadoopCNV with several other software tools using a WGS dataset on a 10-member pedigree generated in house (41). For Illumina sequencing, we quantified the samples using Qubit^®^ dsDNA BR Assay Kit from Invitrogen (Carlsbad, CA, USA), and 1μg of each sample was sent out for WGS using the Illumina^®^ Hiseq 2000 platform. Sequencing libraries were generated from 100ng of genomic DNA using the Illumina TruSeq Nano LT kit, according to manufacturer recommendations. The quality of each library was evaluated with the Agilent bioanalyzer high sensitivity assay (less than 5% primer dimers), and quantified by qPCR (Kappa Biosystem, CT, USA). The pooled library was sequenced in three lanes of a HiSeq2000 paired end 100bp flow cell. The number of clusters passing initial filtering was above 80%, and the number of bases at or above Q30 was above 85%. These samples are used for assessing the Mendelian precision of CNV calls, which is a widely accepted measure of error rates for CNV calling algorithms.

### Implementation of HadoopCNV

Short reads were aligned using BWA-mem (34) to the reference genome, and the BAM file is used as input for HadoopCNV. Processing three billion bases encoded in BAM files poses a significant I/O burden. While it is possible to efficiently subsample a (random or otherwise) fraction of the sites because BAM files are indexed, this suboptimal approach may miss critical sites that are informative for copy number changes. More importantly, this heuristic is likely to miss many of the polymorphic sites that are informative for heterozygosity status. To alleviate the heavy data processing demands, we exploited the benefits of a mature parallel programming framework - Apache Hadoop. Hadoop implements the MapReduce (MR) design in which very large datasets are divided into subsets that are physically stored on multiple distributed data nodes of a cluster. By processing data in a Map stage, bandwidth demands on both shared storage and local area network links are minimized.

After the Map step by Mappers, which can be considered to be “embarrassingly parallel”, data is summarized by a Reduce stage, which entails shuffling of some data across the Mappers running on different compute nodes. In the context of CNV calling, we carry out the following pipeline, which consists of three MR components that execute in the following sequential order:

> **Depth Caller MR:** This component is the most data intensive step, as it entails parsing base-allele specific map and read quality scores for each read across all sites of a whole genome. Mappers in this component make use of the Hadoop-BAM library (42) for extracting a single sequence alignment/map (SAM) record from the BAM files. SAM records are exposed by the Java API of HTSlib, which is distributed as part of SAMtools (43). A Mapper aligns read information within a fixed size view. In practice we set the default to a 100 base pair view, though this is a user-adjustable parameter. Key-value pairs are emitted by the mapper where the key is comprised of the chromosome and the first base pair position of the view. One-byte integer arrays are emitted as values, where the even indices (assuming a 0-based index scheme) store the allele code and the odd indices store their respective phred scores (44) of the allele stored in the preceding array element. Reducers collect all byte arrays that correspond to the chromosome and frame, and output site and allele specific quality weighted depth scores at base pair resolution, where depth score is the sum of the quality scores converted from phred scores: computed as 1 – 10^−0.1*phred_score^.
>
> **Region Binner MR:** The output from the Depth Caller is enormous (can be hundreds of GB) as it contains the cross product of the average number of alleles and sites, where each record is stored as ASCII text on HDFS. The statistics we are most interested in is the median overall depth count and the Hidden Markov Model (HMM) state-specific mean squared errors (SSMSE) conditional on the observed allelic intensity ratios. To compute SSMSE, we first determine which sites within a bin are polymorphic with high confidence based on the user supplied VCF file. Here we adopted the BAF (B Allele Frequency) concept from PennCNV, as the fraction of reads with alternative alleles over all reads; however, to simplify matters, we consider the minor allele as “B” allele so that BAF values are constrained to be in the range of 0 to 0.5. The heterozygous site is determined by the user-provided VCF file. The SSMSE is then calculated by averaging squared errors for each state. SSMSE=BAF^2^ for all sites with homozygous SNVs; otherwise, SSMSE is calculated as
>
> 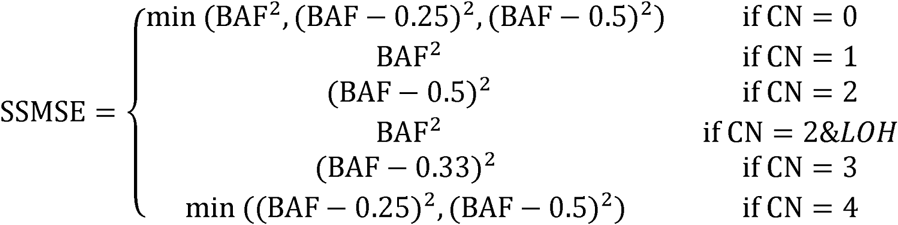
> Output from this component’s Reducer includes start and end location for the bin, median depth score, the SSMSE vector, and the total sites in the bin.
>
> **CNV Caller MR:** At this point we consider the window-specific summary statistics described above as observed data points for CNV inference. For a given chromosome, the Reducer of this component takes all observed summary statistics sorted by position as input, where each window can be represented as each node. The Dynamic Programming Imputation (DPI) method is a generalization of HMMs, which has been successfully applied to CNV inference in microarrays (31). In contrast to a standard HMM where one assumes a probability distribution for each state in the emission matrix, the DPI method loosens this constraint by minimizing losses rather than maximizing a likelihood at each state.
>
> At each node, the median depth is transformed to a *y* value: first, the mean depth of the whole chromosome is calculated as *μ*, and the standard deviation is calculated as *σ*; second, the depth at each node is normalized by deduction of *μ* and division by *σ*; By default, there are a total of 6 states (*s*_0_, *s*_1_, *s*_2_, *s*_3_, *s*_4_, *s*_5_), corresponding to copy number 0, copy number 1, copy number 2 homozygous, copy number 2 heterozygous, copy number 3 and copy number 4 respectively, with expected *y* as -1.0, -0.5, 0, 0, 0.5 and 1.0. The state space can be readily expanded to accommodate more copy numbers.
>
> The loss function is defined as:
>
> 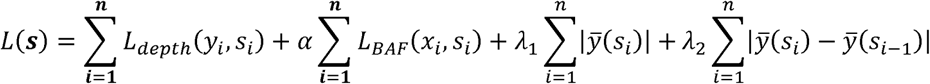
> where *L*_*depth*_(*y_i_*,*S_i_*) is the squared error from the expected *y* in state *i*. *L*_*baf*(*X_i_*,*S_i_*_) is the calculated average SSMSE from previous step for state i, 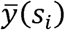 is the expected y for state I, *λ*_1_ and *λ*_2_ are tuning parameters for the penalties, where a large *λ*_1_ penalizes any deviation from copy number 2, and a large *λ*_2_ penalizes a state transition. In practice, is set to 0.01 and *λ*_2_ is set to 1.6. This loss function is minimized by Dynamic Programming starting from the first node to the last, with discrete states at each node. The Viterbi algorithm is used to retrieve the optimized state at each node.
>
> **Post processing:** the output from all the steps above is a copy number and state for each bin for the whole genome. We post-processed them by merging the neighbor deletions or duplications recursively, as long as their distance is less than 20% of their total length, until no two deletions or duplications can be further merged. By default, we filtered out the CNVs less than 500 bp as they are more likely caused by noises, due to the lack of use of split-read or paired-end distance information. Therefore, HadoopCNV is complementary to methods that leverage split-read or paired-end distance information, and their combined use may help reveal the full range of CNVs with different sizes.

### CNV calling on WGS data by other software

CNVnator is used with a 100bp-bin as recommended in its README file, which uses an algorithm called ‘mean-shift’, from image processing, to call CNVs. LUMPY is also used with the recommended pipeline from its document at https://github.com/arq5x/lumpy-sv, which focuses on utilizing information from paired-end reads and split reads to infer break points and CNVs. For LUMPY, the output contains two CNV breakpoint estimates, which are averaged as a single point for the start or end of the CNV.

### Performance evaluation

All the CNV calls from each tool are first compiled into a standard format and then tested with the same comparison script. The CNV calls are first sorted by their physical positions, then the overlapping CNVs are merged into a single CNV. The precision and recall of NA12878 CNVs are calculated with the published gold standard CNV calls as true positive CNVs (37).

Two different standards are used to calculate the overlap: 1) the touch-based standard treats two CNVs as overlap if they share at least one single base pair; 2) the 50% reciprocal standard counts two CNVs as overlap only if their shared length is no less than 50% of their total non-overlapping length. The concordance rate is calculated as the number of overlapping CNVs divided by the total number of CNVs.

### Combination of HadoopCNV and LUMPY result

The script to combine HadoopCNV and LUMPY is written in Perl. First, we determine the concordant CNVs, for deletions and duplications respectively, with the 50% reciprocal overlap standard. The CNV calls from HadoopCNV and LUMPY not in the concordant set are directly included into the combined set. For the concordant ones, all CNVs are included in the final call set, but with the breakpoints calculated from LUMPY.

### Availability

HadoopCNV is available at https://github.com/WGLab/HadoopCNV. Users need to install Hadoop2.0+ on the system, which could be a computing cluster with many nodes or merely a single machine. Therefore, for users without access to large-scale clusters, HadoopCNV can still be executed in single node mode.

## Acknowledgements

This project was supported by NIH grant R01 HG006465. We thank the Wang lab members for helpful comments and suggestions.

## Author Contributions

K.W. conceived and designed the study. G.K.C., H.Y. and M.H. developed the algorithms and implemented software tools. L.L. performed data simulation. H.Y. and L.L performed benchmarking experiments and compared performance with competing tools. H.F. L.J. and G.J.L. provided materials and testing data. G.K.C., H.Y., M.H., and K.W. wrote the manuscript. M.L. guided algorithm development and edited the manuscript. All authors read and approved the manuscript.

